# Protein kinase A controls the hexosamine pathway by tuning the feedback inhibition of GFAT-1

**DOI:** 10.1101/2020.07.23.216424

**Authors:** Sabine Ruegenberg, Felix A.M.C. Mayr, Stephan Miethe, Ilian Atanassov, Ulrich Baumann, Martin S. Denzel

## Abstract

The hexosamine pathway (HP) is a key anabolic pathway whose product uridine 5’-diphospho-N-acetyl-D-glucosamine (UDP-GlcNAc) is an essential precursor for all glycosylation processes in mammals. It modulates the ER stress response, is implicated in cancer and diabetes, and HP activation extends lifespan in *Caenorhabditis elegans*. The highly conserved glutamine fructose-6-phosphate amidotransferase 1 (GFAT-1) is the first and rate-limiting HP enzyme. GFAT-1 activity is modulated through UDP-GlcNAc feedback inhibition and by kinase signaling, including Ser205 phosphorylation by protein kinase A (PKA). The consequence and molecular mechanism of GFAT-1 phosphorylation, however, remains poorly understood. Here, we identify the GFAT-1 R203H substitution that elevates UDP-GlcNAc levels in *C. elegans*, leading to ER stress resistance. In human GFAT-1, the R203H substitution interfered with both UDP-GlcNAc inhibition and PKA-mediated Ser205 phosphorylation. Of note, Ser205 phosphorylation had two discernible effects: It lowered baseline GFAT-1 activity and abolished UDP-GlcNAc feedback inhibition. Thus, GFAT-1 phosphorylation by PKA uncoupled the feedback loop of the HP and, depending on UDP-GlcNAc availability, phosphorylation by PKA results in lower or enhanced GFAT-1 activity *in vivo*. Mechanistically, our data indicate that the relative positioning of the two GFAT-1 domains might be affected by phosphorylation and we propose a model how Ser205 phosphorylation modulates the activity and feedback inhibition of GFAT-1.

## Introduction

The hexosamine pathway (HP) converts fructose-6-phosphate (Frc6P) to uridine 5’-diphospho-N-acetyl-D-glucosamine (UDP-GlcNAc) (Fig. 1a)^1^. 2 to 3 % of cellular glucose enter the HP, as well as L-glutamine (L-Gln), acetylcoenzyme A, and uridine^2^. Hence, the HP integrates sugar, amino acid, fatty acid, and nucleotide metabolism and is considered an important nutrient sensing pathway. The GlcNAc moiety of UDP-GlcNAc is used as a building block for several macromolecules, including peptidoglycans in bacteria, chitin in fungi and insects, or glycosaminoglycans such as hyaluronic acid in vertebrates^3–8^. Moreover, UDP-GlcNAc is an essential precursor for glycosylation reactions in eukaryotes. N-linked glycosylation takes place in the endoplasmic reticulum (ER) and is crucial for proper protein folding and the solubility of proteins^9^. Mucin-type O-linked glycosylation occurs in the Golgi apparatus and is found on many cell surface and secreted proteins^10^. In protein O-GlcNAcylation that occurs in cytoplasm and nucleus, UDP-GlcNAc is the donor for the attachment of a single GlcNAc moiety to serine or threonine residues^11^. This process can fine tune a protein’s function akin to phosphorylation and often O-GlcNAcylation and phosphorylation compete for modification of the same sites^12,13^.

**Fig. 1:**
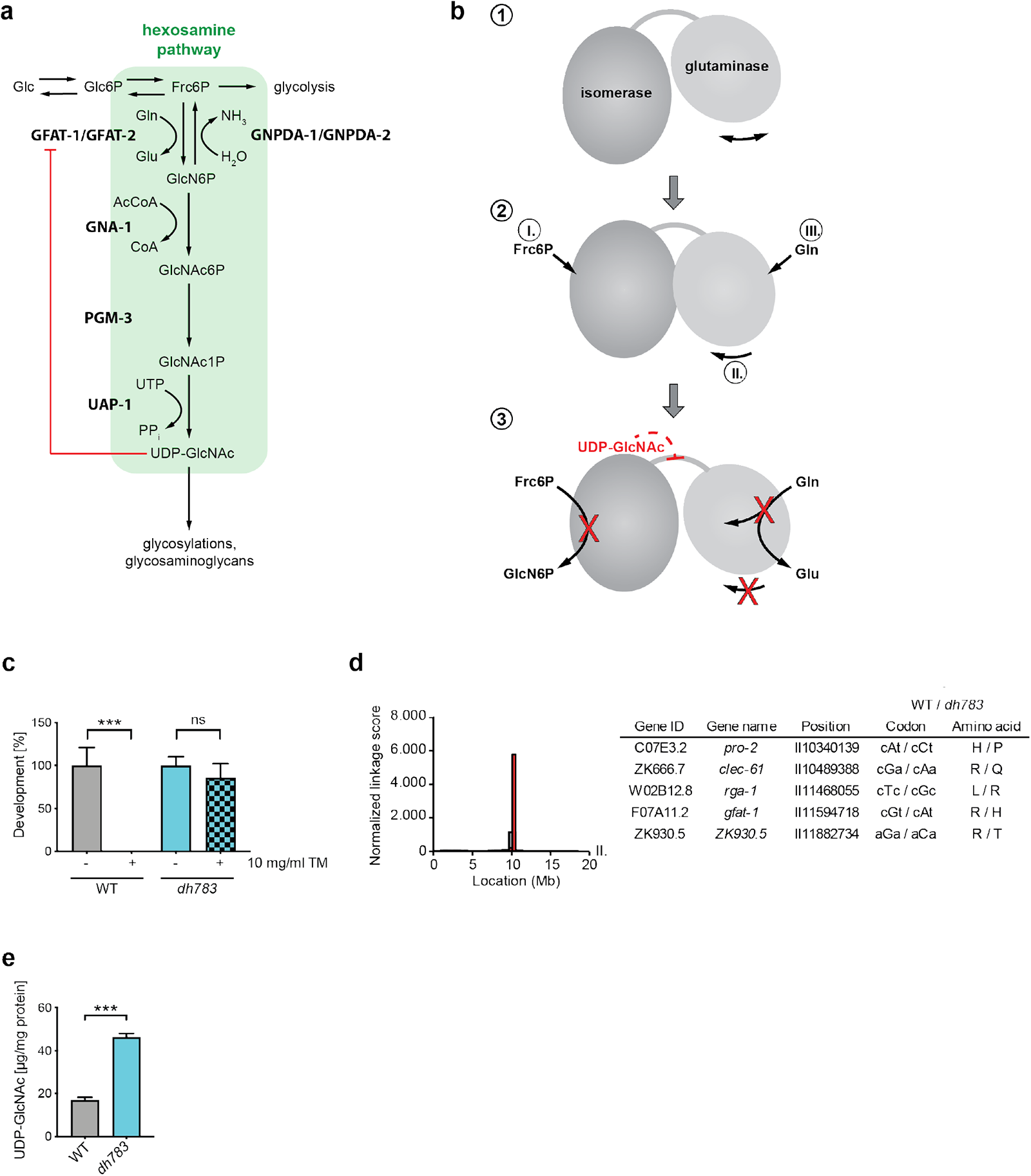
Characterization of *gfat-1(dh783) C. elegans* mutants. **a,** Schematic representation of the hexosamine pathway (green box). The enzymes in the pathway are glutamine fructose-6-phosphate amidotransferase (GFAT-1/-2), glucosamine-6-phosphate N-acetyltransferase (GNA-1), phosphoglucomutase (PGM-3), UDP-N-acetylglucosamine pyrophosphorylase (UAP-1), and glucosamine-6-phosphate deaminase (GNPDA-1/-2). UDP-GlcNAc inhibits eukaryotic GFAT (red line). **b**, Catalytic scheme of one GFAT monomer. (1) Before catalysis: The glutaminase domain does not adopt a fixed position. (2) Substrate binding: I. Frc6P binds, II. the glutaminase domain adopts a specific position, III. L-Gln binds. (3) Catalysis and UDP-GlcNAc inhibition. Catalysis: L-Gln is hydrolyzed to L-Glu and the released ammonia is shuttled through an ammonia channel from the glutaminase to the isomerase domain. There, Frc6P is isomerized to Glc6P and the ammonia is transferred to build GlcN6P. UDP-GlcNAc inhibition: UDP-GlcNAc binds to the isomerase domain, interacts with the interdomain linker, and inhibits the glutaminase function and thereby the GlcN6P production. **c**, *C. elegans* (N2 wild type and *gfat-1*(*dh783*)) developmental resistance assay on NGM plates containing 10 mg/ml tunicamycin. **d**, Frequency plot of normalized parental alleles on chromosome II of *dh783*. The CloudMap Hawaiian Variant Mapping with WGS tool displays regions of linked loci where pure parental allele SNP positions instead of allele positions containing Hawaiian SNPs are over-represented. Gray bars represent 1 Mb and red bars represent 0,5 Mb sized bins. Table: Candidate non-synonymous SNPs between 10 and 12 Mb on chromosome II of *gfat-1*(*dh783*) animals. **e**, UDP-GlcNAc levels in N2 wild type and *gfat-1*(*dh783*) animals (mean +SD, n=5, *** p<0.001, unpaired t-test).

HP activity is regulated by its first and rate-limiting enzyme glutamine fructose-6-phosphate amidotransferase (GFAT, EC 2.6.1.16)^2^. Two GFAT paralogs exist that primarily differ in their tissue-specific expression patterns^14^. GFAT is a modular enzyme composed of a glutaminase domain responsible for hydrolysis of L-Gln into L-glutamate (L-Glu) and an isomerase/transferase domain that catalyzes the isomerization of Fru6P to glucose-6-phosphate (Glc6P) as well as the transfer of ammonia to Glc6P to build glucosamine-6-phosphate (GlcN6P)^15^. The orientation of the two domains with their respective active sites is relevant for catalysis and affected by substrate binding. In the absence of Frc6P, the glutaminase domain is very flexible (Fig. 1b, 1) and no corresponding electron density is visible in the *E. coli* GFAT structure, although SDS-PAGE analyses of dissolved crystals indicate the presence of the full-length protein^16,17^. The reaction is initiated by binding of Frc6P to the isomerase domain, which triggers the glutaminase domain to adopt a specific position relative to the isomerase domain^16^ (Fig. 1b, 2). Subsequently, L-Gln binds to the glutaminase domains, closing the pocket of the glutaminase active site^18^. This is accompanied by a rotation of the glutaminase domain by 21° relative to the isomerase domain^19^ (Fig. 1b, 2). Finally, a solvent-inaccessible channel is formed that links the two active sites to allow the diffusion of ammonia^19–21^.

A disturbed function of GFAT-1 due to mutations within *gfat-1* cause limb-girdle congenital myasthenic syndrome with tubular aggregates^22,23^. This inherited disorder is characterized by defective neuromuscular transmission through impaired neuromuscular junctions, which transmit impulses from motor neurons to skeletal muscle fibers^22^. Moreover, many studies suggest a role of the HP and GFAT in the development and progression of diabetes or cancer^24–26^. Both disorders are characterized by an abberant glucose metabolism, and the HP links altered metabolism with aberrant glycosylation. Especially elevated O-GlcNAcylation contributes to the pathogenesis of diabetes and cancer. Altered O-GlcNAcylation was also reported to play a critical role in neurodegenerative diseases, heart disease, and inflammation^27,28^.

Given that GFAT drives the metabolic flux of the HP, its regulation is of great physiological importance and in eukaryotes it occurs through UDP-GlcNAc feedback inhibition^29–31^ and through phosphorylation events. In human GFAT-1, Ser205 and Ser235 are phosphorylated by cAMP-dependent protein kinase (PKA)^32–34^. Ser205 phosphorylation has reported effects on GFAT-1 activity, although the published data are contradictory: On the one hand, phosphorylation of Ser205, and the corresponding Ser202 in human GFAT-2, was reported to increase activity^32,34^. On the other hand, it was reported that Ser205 phosphorylation leads to decreased GFAT-1 activity^33^. Phosphorylation of the second PKA site Ser235 in GFAT-1, which is not conserved in all eukaryotes and is absent in GFAT-2, seems not to affect the enzymatic activity^33,35^. Ser243 was shown to be phosphorylated by adenosine monophosphate (AMP)-activated protein kinase (AMPK) and calcium/calmodulin-dependent kinase II, but their effect on GFAT activity remains elusive. Phosphorylation at this site is reported to be activating or inhibiting^36–38^.

Previously, in a *C. elegans* forward genetic mutagenesis screen we identified gain-of-function mutations in *gfat-1* that suppressed tunicamycin-induced proteotoxic stress^39^. These mutations increased UDP-GlcNAc concentrations in the worms that also showed improved protein quality control as well as a significant lifespan extension^39^. Recently, we showed that loss of regulation by UDP-GlcNAc feedback inhibition constitutes a gain-of-function mechanism^40^. Although UDP-GlcNAc binds to the isomerase domain of GFAT-1, this binding inhibits the glutaminase function and consequently the GlcN6P production of the whole enzyme^41^. We identified a critical role of the interdomain linker during UDP-GlcNAc inhibition and proposed that UDP-GlcNAc disturbs the tight coupling of the active sites by interfering with the relative orientation of the two domains^40^ (Fig. 1b, 3).

Here, we identify the R203H GFAT-1 gain-of-function mutation that interferes with UDP-GlcNAc inhibition and with PKA-dependent phosphorylation at Ser205. Analyses of the phospho-mimic S205D substitution resolved the controversially discussed effect of Ser205 phosphorylation. Our data demonstrate that PKA phosphorylation at Ser205 lowers GFAT-1 activity and simultaneously blocks UDP-GlcNAc inhibition. We propose a model how Ser205 phosphorylation might modulate the activity and feedback inhibition of GFAT-1 through phosphorylation-induced domain movement.

## Results

### A new GFAT-1 gain-of-function point mutant is resistant to ER stress

Previously, we performed a forward genetic screen in *C. elegans* for mutants that are resistant to tunicamycin-induced proteotoxic stress^39^. From this screen, we obtained the mutant allele *dh783*, which was not characterized until now. While 10 mg/ml tunicamycin was toxic for N2 wild type worms, the *dh783* allele conferred strong tunicamycin resistance (Fig. 1c, Supplementary Fig. 1a). To identify the causal mutation, we performed Hawaiian single nucleotide polymorphism (SNP) mapping^42^. The normalized linkage score identified SNPs in exons of five candidate genes on chromosome II including *gfat-1* (Fig. 1d). Given that tunicamycin toxicity can be suppressed by elevated UDP-GlcNAc levels through gain-of-function mutations in *gfat-1*^39^, we quantified UDP-GlcNAc and its epimer UDP-GalNAc by mass spectrometry. Indeed, we found significantly elevated UDP-GlcNAc and UDP-GalNAc levels in whole worm lysates in mutant carriers of the *dh783* allele (Fig. 1e, Supplementary Fig. 1b). We conclude that tunicamycin resistance was caused by the *gfat-1*(*dh783*) allele due to the GFAT-1 R203H gain-of-function substitution.

### The GFAT-1 R203H substitution interferes with UDP-GlcNAc inhibition leading to gain-of-function

GFAT-1 is well-conserved from *C. elegans* to humans (Supplementary Fig. 2a). To decipher the effect of the R203H substitution, we crystallized human GFAT-1 R203H and characterized it in activity assays. GFAT-1 R203H crystals diffracted to a resolution limit of 2.77 Å (Table 1). Overall, compared to wild type GFAT-1, the R203H substitution does not cause major structural changes (Fig. 2a, b, Supplementary Fig. 2b). While Arg203 interacts with the backbone residues of Ile178 and Leu181, as well as with Asp279 in wild type GFAT-1, these interactions are abolished in the R203H GFAT-1 (Fig. 2b). In the mutant enzyme, however, the side chain of His203 protrudes between Asp278 and Asp279, thereby stabilizing the orientation of the neighboring loop (residues 277-280) in a similar position as in the wild type (Fig. 2b). Subsequently, we analyzed the R203H substitution of human GFAT-1 in activity assays and found similar kinetics for L-Glu and GlcN6P production as in wild type GFAT-1 (Fig. 2c, d, Table 2). In contrast, UDP-GlcNAc dose response assays revealed an approximately 6-fold higher IC_50_ value of R203H GFAT-1 compared to wild type GFAT-1 (R203H: 231.7 −15.2/+16.3 μM; wild type: 41.4 −3.5/+3.9 μM) (Fig. 2e). This finding suggests that a lower sensitivity to UDP-GlcNAc feedback inhibition leads to the gain-of-function caused by the R203H substitution *in vivo*.

**Fig. 2:**
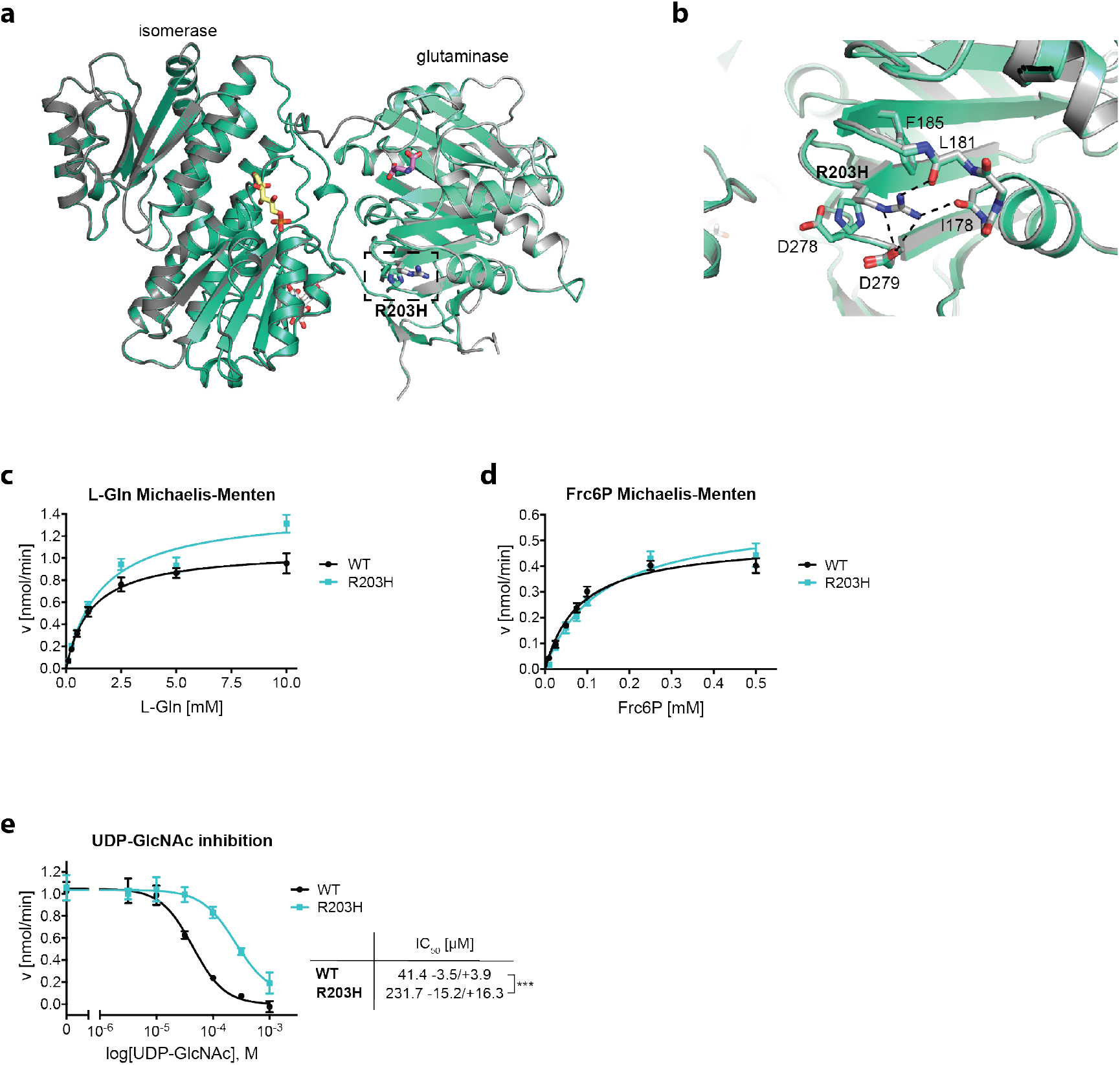
The GFAT-1 R203H gain-of-function substitution perturbs UDP-GlcNAc feedback inhibition. **a, b,** Position of the R203H mutation in the structure of GFAT-1. Proteins are presented as cartoons. Superposition of wild type GFAT-1 (light gray/dark gray) and R203H GFAT-1 (green-cyan, teal). Glc6P (yellow sticks), L-Glu (violet sticks), and UDP-GlcNAc (white sticks) are highlighted, as well as the position of R203H (black box). **a**, Overview. R203H (sticks) is located at the glutaminase domain of GFAT-1 (dashed box). **b**, Close-up view of the position of R203H focusing on residues in close proximity. The R203H mutation and residues in close proximity to the mutation are highlighted with sticks. Arg203 interacts with the neighboring loops (dashed lines). **c**, L-Gln kinetic of wild type (WT, black circle) and R203H (cyan square) GFAT-1 (mean ± SEM, WT n=5, R203H n=4). **d**, Frc6P kinetic of wild type (black circle) and R203H (cyan square) GFAT-1 (mean ± SEM, WT n=5, R203H n=4). **e**, Representative UDP-GlcNAc dose response assay of wild type (black circle) and R203H (cyan square) GFAT-1 (mean ±SD, n=3). Table: IC_50_ UDP-GlcNAc values (mean ±SEM, n=4, ***p<0.001, unpaired t-test).

**Table 1.**
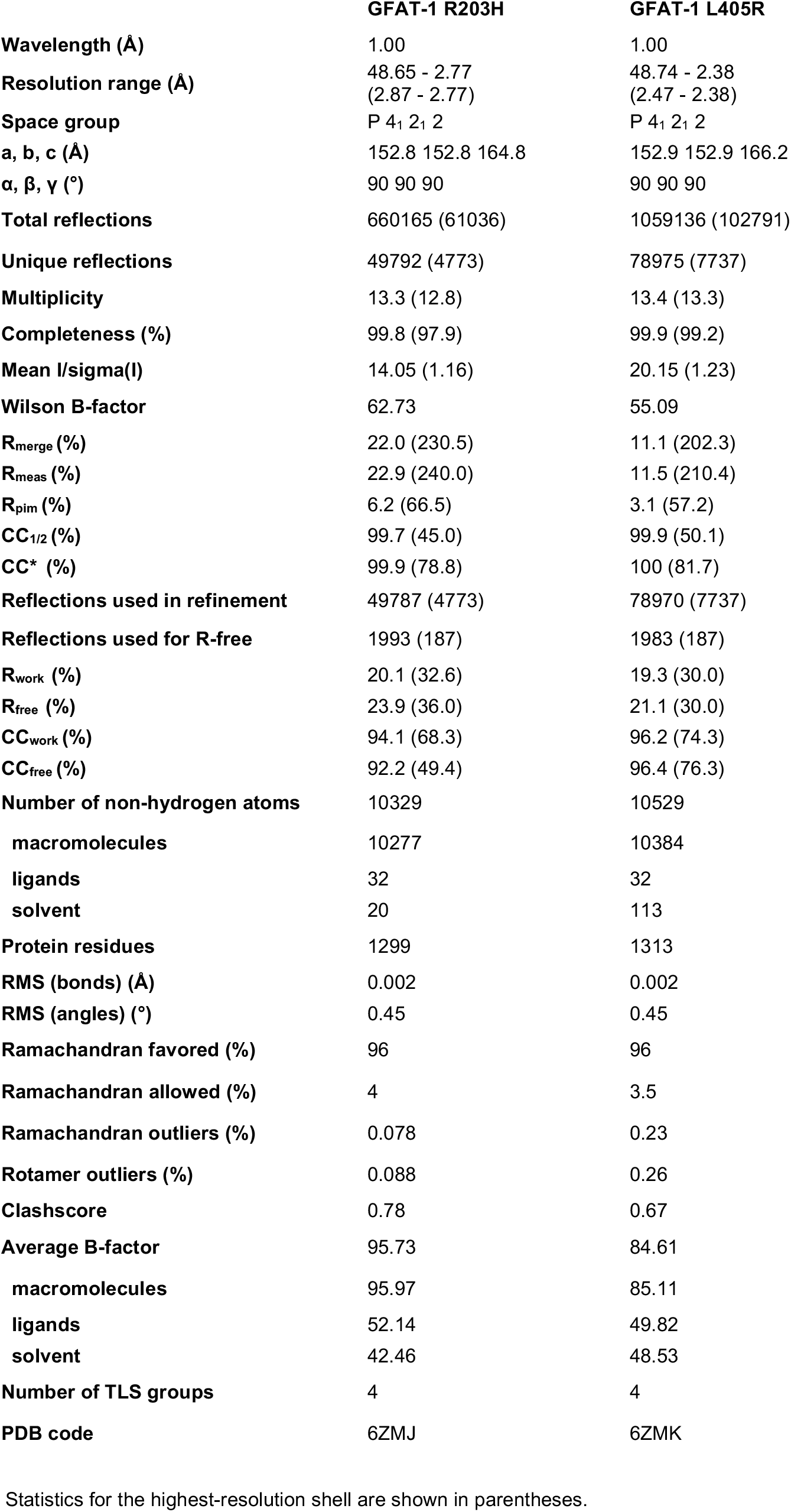
Data collection and refinement statistics.

|  | **GFAT-1 R203H** | **GFAT-1 L405R** |
| --- | --- | --- |
| **Wavelength (Å)** | 1.00 | 1.00 |
| **Resolution range (Å)** | 48.65 - 2.77<br>(2.87 - 2.77) | 48.74 - 2.38<br>(2.47 - 2.38) |
| **Space group** | P 4<sub>1</sub> 2<sub>1</sub> 2 | P 4<sub>1</sub> 2<sub>1</sub> 2 |
| **a, b, c (Å)** | 152.8 152.8 164.8 | 152.9 152.9 166.2 |
| **α, β, γ (°)** | 90 90 90 | 90 90 90 |
| **Total reflections** | 660165 (61036) | 1059136 (102791) |
| **Unique reflections** | 49792 (4773) | 78975 (7737) |
| **Multiplicity** | 13.3 (12.8) | 13.4 (13.3) |
| **Completeness (%)** | 99.8 (97.9) | 99.9 (99.2) |
| **Mean I/sigma(I)** | 14.05 (1.16) | 20.15 (1.23) |
| **Wilson B-factor** | 62.73 | 55.09 |
| **R<sub>merge</sub> (%)** | 22.0 (230.5) | 11.1 (202.3) |
| **R<sub>meas</sub> (%)** | 22.9 (240.0) | 11.5 (210.4) |
| **R<sub>pim</sub> (%)** | 6.2 (66.5) | 3.1 (57.2) |
| **CC<sub>1/2</sub> (%)** | 99.7 (45.0) | 99.9 (50.1) |
| **CC\* (%)** | 99.9 (78.8) | 100 (81.7) |
| **Reflections used in refinement** | 49787 (4773) | 78970 (7737) |
| **Reflections used for R-free** | 1993 (187) | 1983 (187) |
| **R<sub>work</sub> (%)** | 20.1 (32.6) | 19.3 (30.0) |
| **R<sub>free</sub> (%)** | 23.9 (36.0) | 21.1 (30.0) |
| **CC<sub>work</sub> (%)** | 94.1 (68.3) | 96.2 (74.3) |
| **CC<sub>free</sub> (%)** | 92.2 (49.4) | 96.4 (76.3) |
| **Number of non-hydrogen atoms** | 10329 | 10529 |
| **macromolecules** | 10277 | 10384 |
| **ligands** | 32 | 32 |
| **solvent** | 20 | 113 |
| **Protein residues** | 1299 | 1313 |
| **RMS (bonds) (Å)** | 0.002 | 0.002 |
| **RMS (angles) (°)** | 0.45 | 0.45 |
| **Ramachandran favored (%)** | 96 | 96 |
| **Ramachandran allowed (%)** | 4 | 3.5 |
| **Ramachandran outliers (%)** | 0.078 | 0.23 |
| **Rotamer outliers (%)** | 0.088 | 0.26 |
| **Clashscore** | 0.78 | 0.67 |
| **Average B-factor** | 95.73 | 84.61 |
| **macromolecules** | 95.97 | 85.11 |
| **ligands** | 52.14 | 49.82 |
| **solvent** | 42.46 | 48.53 |
| **Number of TLS groups** | 4 | 4 |
| **PDB code** | 6ZMJ | 6ZMK |

Statistics for the highest-resolution shell are shown in parentheses.

**Table 2.**
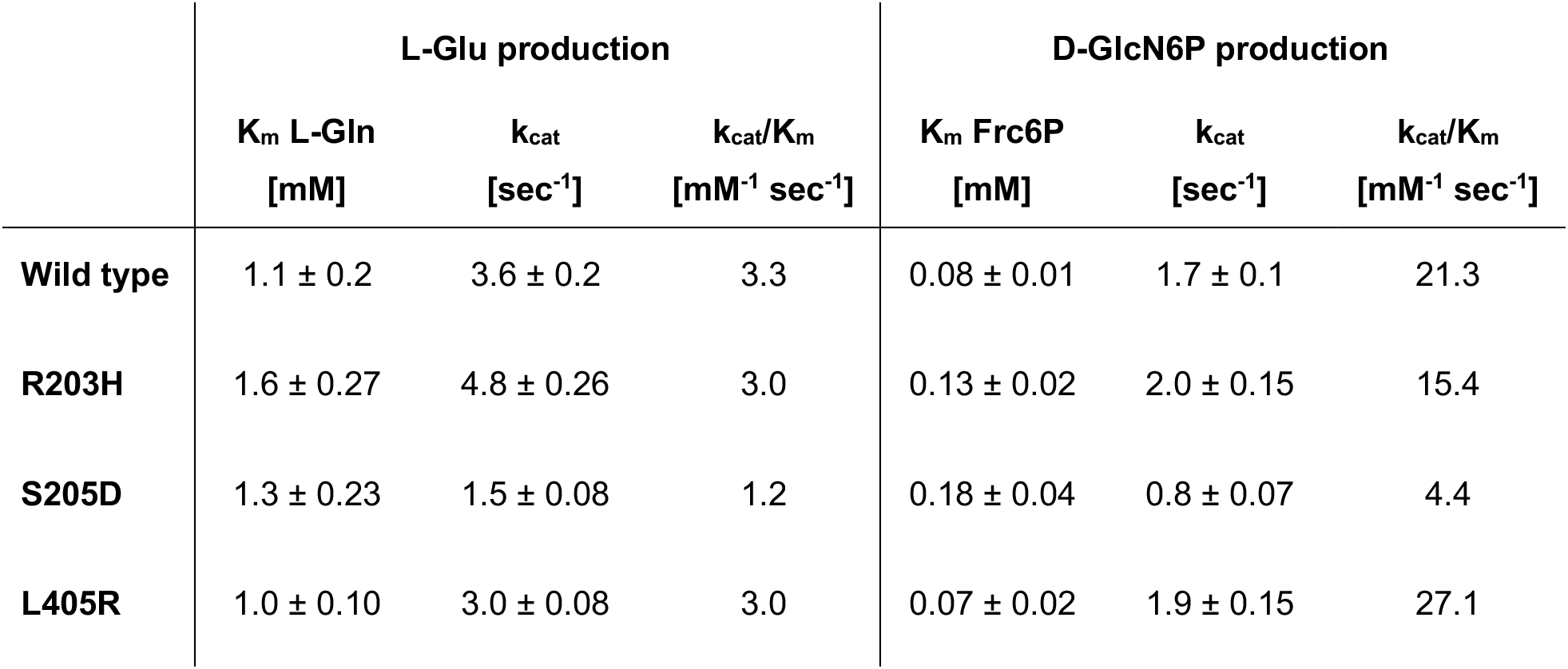
Kinetic parameters.

|  | L-Glu production |  |  | D-GlcN6P production |  |  |
| --- | --- | --- | --- | --- | --- | --- |
| | $K_m$ L-Gln<br>[mM] | $k_{cat}$<br>[sec<sup>-1</sup>] | $k_{cat}/K_m$<br>[mM<sup>-1</sup> sec<sup>-1</sup>] | $K_m$ Frc6P<br>[mM] | $k_{cat}$<br>[sec<sup>-1</sup>] | $k_{cat}/K_m$<br>[mM<sup>-1</sup> sec<sup>-1</sup>] |
| **Wild type** | 1.1 ± 0.2 | 3.6 ± 0.2 | 3.3 | 0.08 ± 0.01 | 1.7 ± 0.1 | 21.3 |
| **R203H** | 1.6 ± 0.27 | 4.8 ± 0.26 | 3.0 | 0.13 ± 0.02 | 2.0 ± 0.15 | 15.4 |
| **S205D** | 1.3 ± 0.23 | 1.5 ± 0.08 | 1.2 | 0.18 ± 0.04 | 0.8 ± 0.07 | 4.4 |
| **L405R** | 1.0 ± 0.10 | 3.0 ± 0.08 | 3.0 | 0.07 ± 0.02 | 1.9 ± 0.15 | 27.1 |

### The GFAT-1 R203H substitution interferes with PKA phosphorylation at Ser205

In addition to the observed gain-of-function caused by reduced feedback inhibition, the R203H substitution alters the PKA consensus sequence of Ser205 from RRGS to RHGS, suggesting a potential change of Ser205 phosphorylation and a subsequent loss of regulation (Fig. 3a). To test whether GFAT-1 R203H can be phosphorylated at Ser205, we purified wild type and R203H GFAT-1 and performed *in vitro* phosphorylation with PKA, followed by LysC protease digest and untargeted as well as targeted mass spectrometric analyses (Fig. 3b). In the untargeted analysis, we identified a number of phosphorylated sites, among them the PKA phosphorylation sites Ser205 and Ser 235^32,33^ (Supplementary Table 1). For the targeted analysis, heavy isotope labeled phosphorylated peptides (Fig. 3b) of wild type or R203H GFAT-1 were spiked in, allowing more sensitive relative quantification of the Ser205 phosphorylation. Untreated insect cell-derived wild type and R203H GFAT-1 preparations showed low detectable levels of phosphorylation at Ser205. After *in vitro* PKA treatment, phosphorylation of wild type GFAT-1 Ser205 was highly induced (Fig. 3c, d). In contrast, the increase of Ser205 phosphorylation was drastically reduced in the presence of the R203H substitution (Fig. 3c, d). The second known PKA phosphorylation site of GFAT-1 at Ser235^33^ was used as a positive control. For this site, we found a significant increase in the abundance of phosphorylated peptides corresponding to Ser235 in both wild type and R203H GFAT-1, proving full activity of PKA in our setup (Supplementary Fig. 3). Thus, disruption of the PKA consensus sequence by the R203H substitution prevents PKA-mediated phosphorylation at Ser205. Taken together, in addition to a lower sensitivity to UDP-GlcNAc inhibition, the GFAT-1 R203H substitution revealed a second gain-of-function mechanism: a loss of PKA-dependent regulation through strongly reduced phosphorylation at Ser205.

**Fig. 3:**
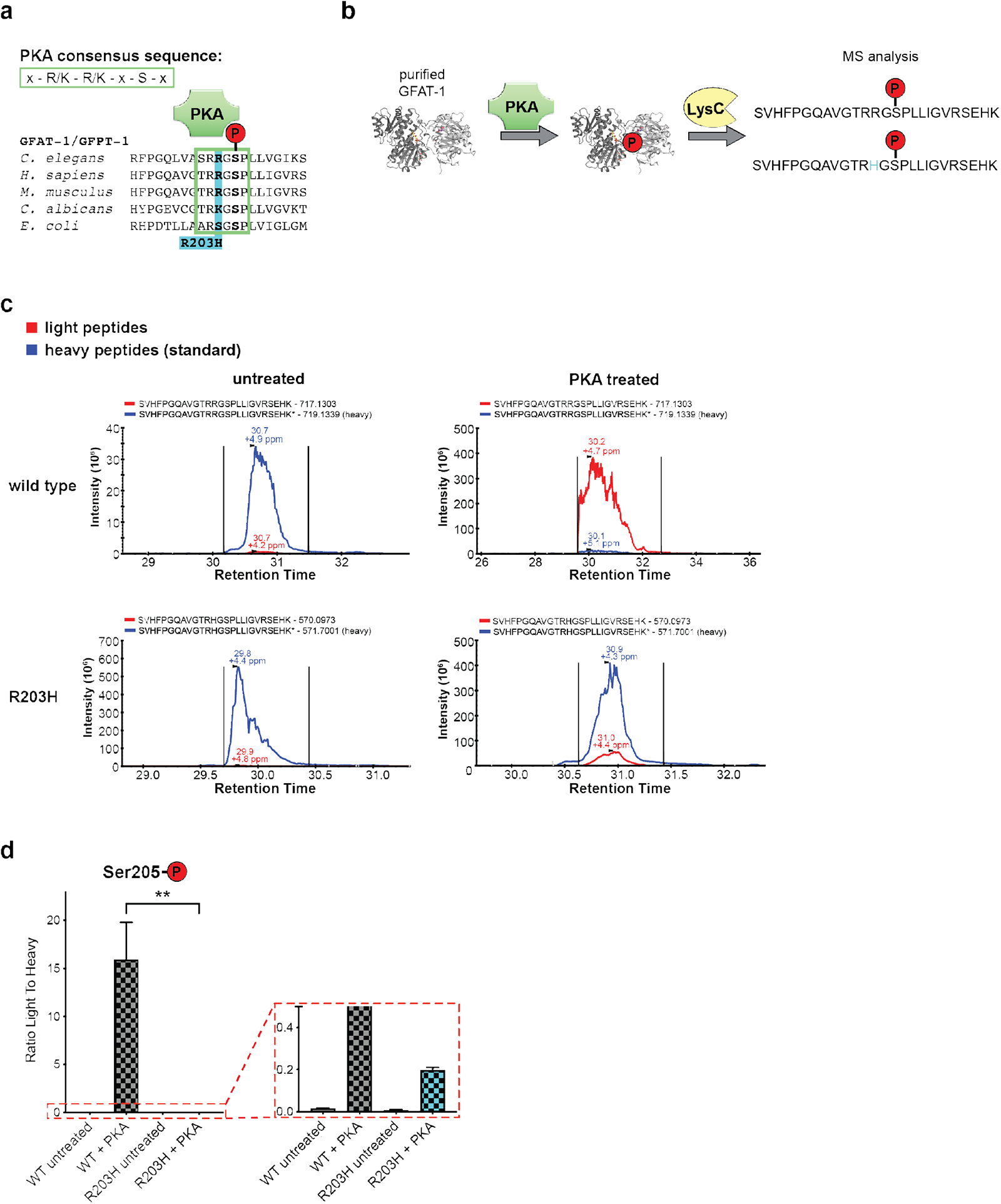
The GFAT-1 gain-of-function substitution R203H disturbs PKA phosphorylation at Ser205. **a,** Position of gain-of-function mutation R203H in a protein sequence alignment of GFAT-1. Mutation R203H (cyan) disrupts the PKA consensus sequence at Ser205. **b**, Workflow for the *in vitro* phosphorylation analysis. **c**, Representative quantification of the light and heavy peptides by mass spectrometry. **d**, Quantification of the Ser205 phosphorylation of wild type GFAT-1 (gray) and R203H (cyan) before and after treatment with PKA (mean +SEM, n=4, ** p<0.01, one-way ANOVA).

### PKA phosphorylation at Ser205 modulates UDP-GlcNAc inhibition of GFAT-1

The loss of PKA phosphorylation in the GFAT-1 gain-of-function R203H variant suggests that the Ser205 phosphorylation might be inhibitory. Until now, the effect of phosphorylation at Ser205 has been controversially reported as either activating^32^ or inhibiting^33^. We aimed to mechanistically understand the effect of Ser205 phosphorylation and generated a GFAT-1 S205D mutant to mimic the phosphorylation. Importantly, wild type insect cell-derived GFAT-1 showed almost no phosphorylation at Ser205 (Fig. 3c, d), permitting the direct comparison with the S205D variant in activity assays. We observed reduced kcat values for L-Glu and GlcN6P production resulting in lower catalytic efficiency (kcat/Km) for both active sites (Fig. 4a, b, Table 2). Overall GlcN6P synthesis was decreased approximately 5-fold in the phospho-mimic mutant (kcat/Km: 4.4 mM^-1^ sec^-1^) compared to wild type GFAT-1 (kcat/Km: 21.3 mM^-1^ sec^-1^) (Table 2). Strikingly, the dose-dependent inhibition by UDP-GlcNAc was completely abolished in the S205D mutant (Fig. 4c). Taken together, these data suggest that PKA-dependent phosphorylation at Ser205 reduces GFAT-1 activity *in vitro*. The additional loss of UDP-GlcNAc feedback inhibition, however, maintains the activity of Ser205-phosphorylated GFAT-1 when UDP-GlcNAc concentrations are high. At low UDP-GlcNAc concentrations up to approximately 30 μM, the phosphorylation is inhibiting, while at high UDP-GlcNAc concentrations above approximately 30 μM, the phosphorylation is activating.

**Fig. 4:**
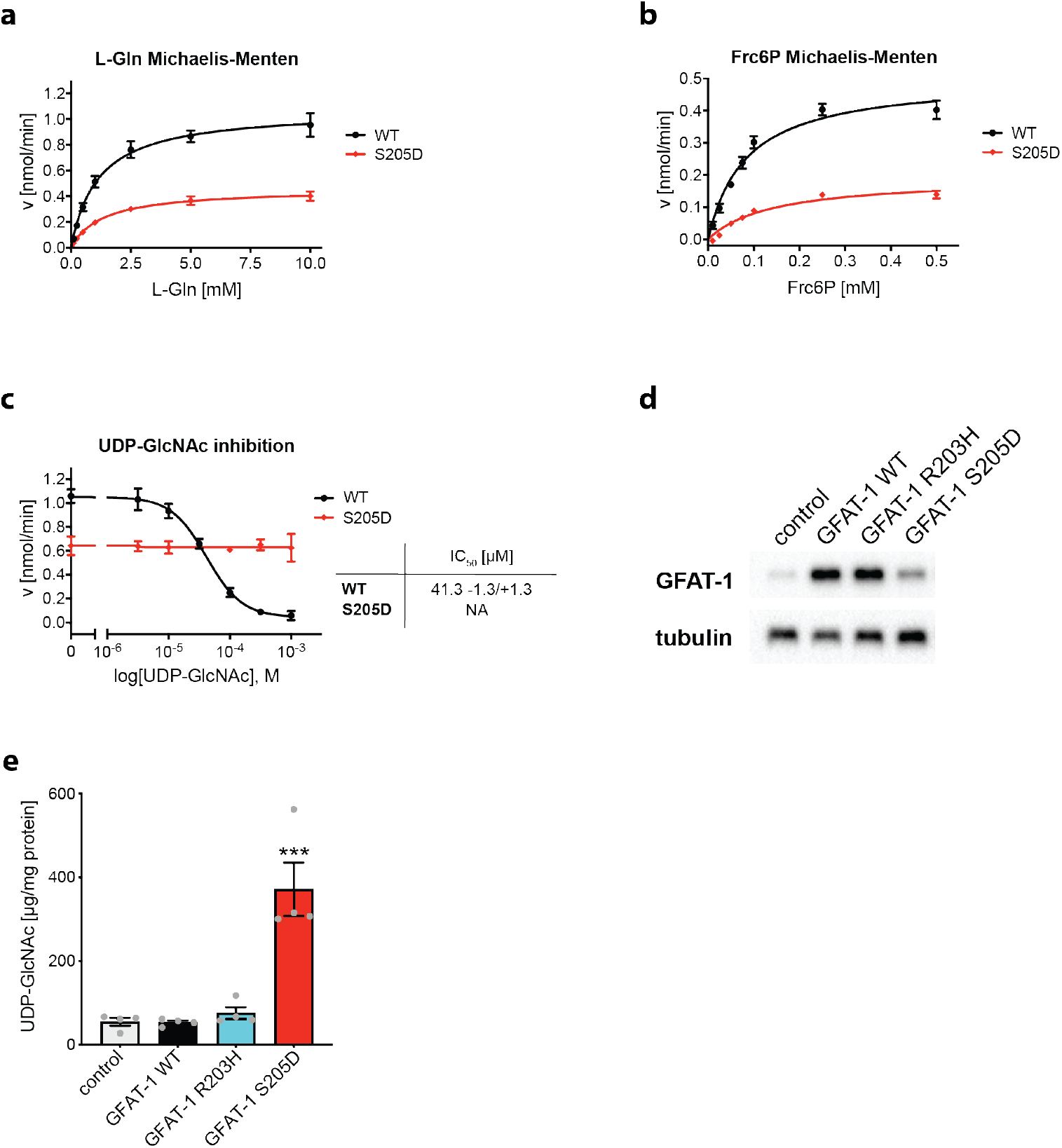
PKA phosphorylation at Ser205 modulates UDP-GlcNAc inhibition of GFAT-1. **a**, L-Gln kinetic of wild type (black circles) and S205D (red diamonds) GFAT-1 (mean ±SEM, WT n=5, S205D n=6). **b**, Frc6P kinetic of wild type (black circles) and S205D (red diamonds) GFAT-1 (mean ±SEM, WT n=5, S205D n=4). **c**, Representative UDP-GlcNAc inhibition of wild type (black circles) and S205D (red diamonds) GFAT-1 (mean ±SD, n=3). Table: IC_50_ UDP-GlcNAc values (mean ±SEM, n=3). **d**, Western blot analysis of GFAT-1 protein levels in control HEK293 cells and HEK293s cell stably overexpressing the indicated GFAT-1 variants. **e**, LC/MS measurement of UDP-GlcNAc normalized to protein content presented as means +SEM with n=4, *** p < 0.001 versus WT, one-way ANOVA.

Understanding the effect of Ser205 PKA phosphorylation on GFAT-1 activity, we next analyzed the relevance of the phosphorylation *in vivo*. For that purpose, we generated HEK293 cell lines, which stably overexpress either wild type, R203H, or S205D GFAT-1 with an internal His6-tag (Fig. 4d) and measured UDP-GlcNAc and UDP-GalNAc levels in the cell lysates (Fig. 4e, Supplementary Fig. 4). The cells were cultured at high glucose concentrations of 25 mM, to avoid any glucose limitation. Strikingly, GFAT-1 overexpression did not elevate UDP-GlcNAc or UDP-GalNAc concentrations. Likewise, the R203H GFAT-1 cell line did not show increased UDP-GlcNAc or UDP-GalNAc levels. This suggests that UDP-GlcNAc feedback inhibition is effective in fully suppressing the activity of wild type and R203H GFAT-1 even during overexpression (Fig. 4e, Supplementary Fig. 4). In contrast, we found significantly elevated levels of UDP-GlcNAc and UDP-GalNAc when the phospho-mimetic S205D variant of GFAT-1 was overexpressed (Fig. 4e, Supplementary Fig. 4). These data support an interference of the S205D substitution with the UDP-GlcNAc inhibition, which rendered GFAT-1 constitutively active. In summary, our data suggest that PKA controls GFAT-1 activity by interference with its UDP-GlcNAc feedback inhibition.

### PKA phosphorylation at Ser205 modulates GFAT-1 domain stability

Next, we assessed the thermal stability of GFAT-1 in thermofluor assays. Full-length GFAT-1 showed two clearly distinguishable melting points at 53.0 °C and 65.0 °C, which are termed “low” and “high” hereafter (Fig. 5a, Supplementary Fig. 5a). Most likely, these two melting points represent the two domains, which possess different stabilities: the isolated isomerase domain melted at 64.0 °C suggesting that the high melting point corresponded to the isomerase domain, while the low melting point was assigned to the glutaminase domain (Fig. 5a, Supplementary Fig. 5a). To investigate their relative interactions, rising salt concentrations were then used to destabilize the domains. For full-length GFAT-1, high salt (0.5 - 1.0 M) lowered the melting point of the glutaminase domain by 3.0-4.0 °C, and it lowered the melting point of the isomerase domain by 1.7-2.0 °C (Fig. 5b). In contrast, the isolated isomerase domain showed a much stronger reduction of the melting temperature of up to −10 °C (Fig. 5b). Together, these results show that in full-length GFAT-1 the isomerase domain is stabilized by the glutaminase domain, thus indicating interdomain interactions. We conclude that analysis of the melting points in GFAT-1 variants can reveal changes in the interaction between its domains. To independently test whether thermofluor data can indeed indicate interdomain interactions, we generated the C-tail lock mutant L405R of human GFAT-1 and characterized it structurally, as well as in thermofluor assays and in kinetic measurements. The well-conserved C-tail (C-terminal residues 670-681) is the major mobile element between the isomerase and glutaminase domain of GFAT. It bears residues of both active sites and mediates interactions between glutaminase and isomerase domain^19^. In *E. coli*, its flexibility is restricted by a salt-bridge between Arg331 and Glu608^43^. Miszkiel and Wojciechowski showed in molecular dynamic simulations that the C-tail lock mutation, which introduces a salt-bridge similar as observed in *E. coli*, limits the C-tail’s flexibility^43^. We solved the crystal structure (resolution 2.38 Å, Table 1) and confirmed that the L405R substitution indeed formed a salt-bridge with the C-terminal Glu681 and additionally interacted with Asp428 (Supplementary Fig. 5b-d). The kinetics (L-Gln and Frc6P) and the UDP-GlcNAc inhibition remained unaffected by the C-tail lock mutation (Table 2, Supplementary Fig. 5e-g). Importantly, the L405R mutant clearly showed a single melting point at 56.9 ± 0.1 °C in thermofluor assays (Supplementary Fig. 5h, i), indicating a changed stability of both domains, which can only be explained by the altered interactions between the two domains. Therefore, thermofluor assays are a valuable tool to analyze interdomain interactions of GFAT-1.

**Fig. 5:**
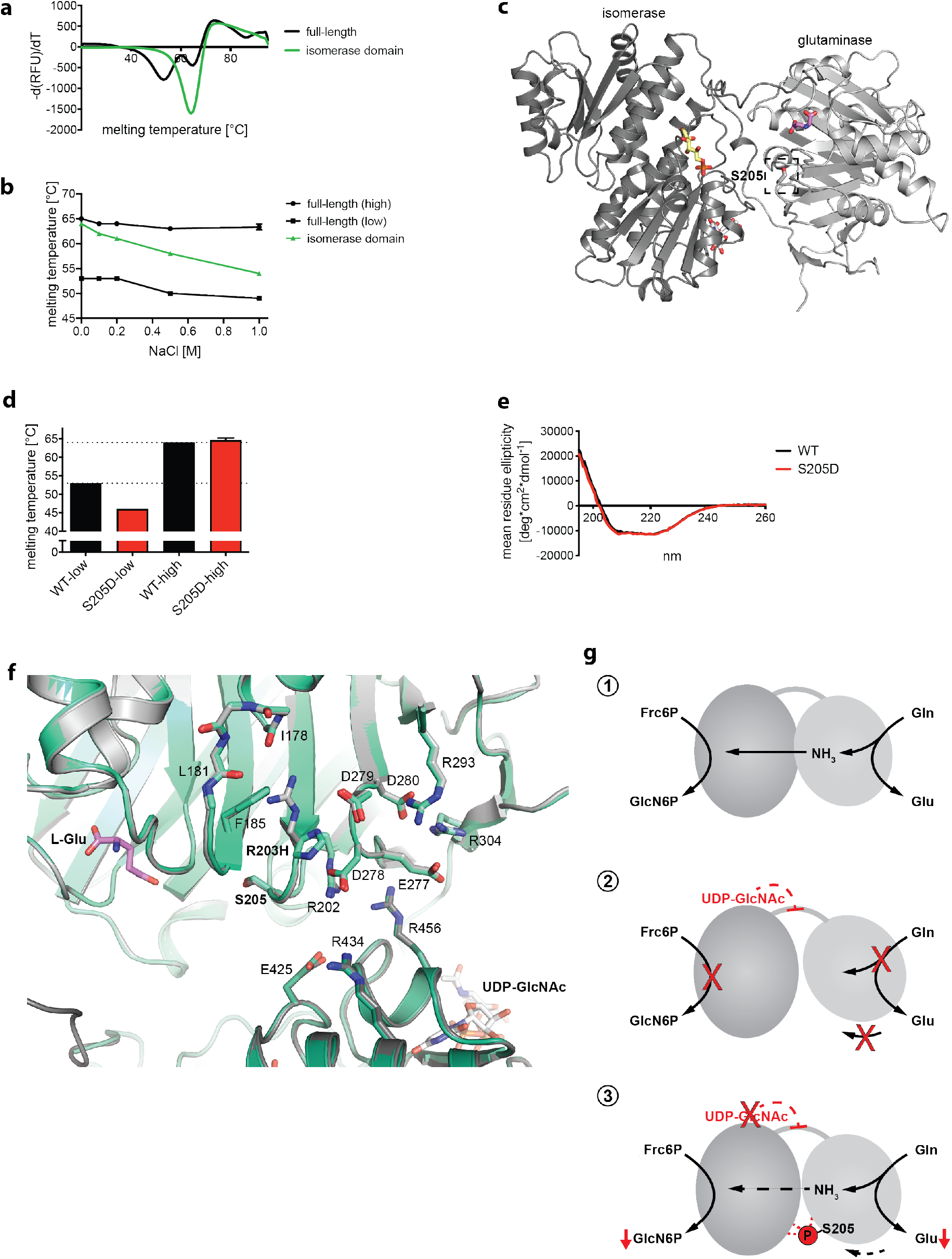
Altered domain stability of GFAT-1 after PKA phosphorylation at Ser205. **a**, Representative derivative melting curves of full-length GFAT-1 (black) and the isolated GFAT-1 isomerase domain (green) in standard SEC buffer without NaCl. **b**, Melting temperatures of full-length GFAT-1 (black) and the isolated GFAT-1 isomerase domain (green) GFAT-1 in SEC buffer with rising NaCl concentrations (mean +SD, n=3). **c**, Overview of the position of the phosphorylation site Ser205 in GFAT-1 in cartoon representation. Glc6P (yellow sticks), L-Glu (violet sticks), and UDP-GlcNAc (white sticks) are highlighted, as well as the PKA site Ser205 (dashed box). **d**, Melting temperatures of wild type GFAT-1 (black) and GFAT-1 S205D (red) (mean +SD, n=3). **e**, CD spectra of wild type GFAT-1 (black) and GFAT-1 S205D (red). **f**, Superposition of the structures of wild type (gray) and R203H (green cyan) GFAT-1 in cartoon representation. L-Glu (violet sticks) and UDP-GlcNAc (white sticks) are highlighted. Ser205, the R203H substitution and residues that might form salt bridges in close proximity to the mutation are highlighted with sticks. **g**, Schematic models of one GFAT monomer. (1) Catalysis: L-Gln is hydrolyzed to L-Glu and the released ammonia is shuttled through an ammonia channel from the glutaminase to the isomerase domain. (2) UDP-GlcNAc inhibition: UDP-GlcNAc binds to the isomerase domain, interacts with the interdomain linker, and inhibits the glutaminase function and thereby the GlcN6P production. (3) Ser205 phosphorylation: Upon phosphorylation, the catalytic activity is reduced and the UDP-GlcNAc inhibition is abolished.

We next used the thermofluor assay to test the role of the phospho-mimic S205D substitution regarding the interdomain interaction of GFAT-1. At physiological pH, phosphorylation introduces a di-anionic phosphate group that causes a repulsion with other negative charges and enables the formation of electrostatic interactions with positively charged residues. The PKA phosphosite Ser205 is located in a loop of the glutaminase domain pointing towards the glutaminase active site and the interdomain cleft between the glutaminase and isomerase domains (Fig. 5c). Ser205 is thus optimally positioned to induce a potential domain movement upon phosphorylation. Notably, GFAT-1 S205D did not form any crystals in the standard GFAT-1 crystallization condition, indicating a structural change compared to wild type GFAT-1 that prevented crystal packing. Compared to wild type GFAT-1, the S205D mutant showed strongly reduced thermal stability of the glutaminase domain, as indicated by the specific reduction of the lower melting point (Fig. 5d, Supplementary 5j, k). The stability of the isomerase domain (high melting point) was not affected (Fig. 5d, Supplementary 5j, k). To rule out major structural changes in GFAT-1 S205D, we performed circular dichroism (CD) measurements of wild type and S205D GFAT-1. Both proteins showed very similar spectra indicating proper folding (Fig. 5e). Together, these data demonstrate a destabilization of the glutaminase domain in the GFAT-1 S205D variant, suggesting altered interdomain interactions through changed domain orientation. Thus, PKA phosphorylation at Ser205 might affect GFAT-1 activity and block the UDP-GlcNAc inhibition through phosphorylation-induced domain movement.

## Discussion

Here we present the GFAT-1 mutant R203H and decipher its two gain-of-function mechanisms: (1) strongly reduced PKA-dependent phosphorylation at Ser205 due to a disrupted consensus sequence and (2) a weaker sensitivity to UDP-GlcNAc feedback inhibition. The current study resolves the contradicting reports regarding the effect of PKA phosphorylation at Ser205 in GFAT-1 and provides a model for phosphorylation-induced domain movement. Strikingly, the phospho-mimetic S205D substitution lowered GFAT-1 activity and abolished UDP-GlcNAc inhibition. Importantly, we demonstrate a modulation of the UDP-GlcNAc feedback inhibition of GFAT-1 by PKA.

We describe two independent gain-of-function mechanisms of GFAT-1 R203H. First, when UDP-GlcNAc levels are low, the lack of phosphorylation at Ser205 results in higher GFAT-1 activity when compared to wild type GFAT-1 with the latter having a reduced GlcN6P production upon phosphorylation. Second, when UDP-GlcNAc levels are higher, the reduced sensitivity to feedback inhibition in R203H GFAT-1 elevates cellular UDP-GlcNAc levels. The observation of elevated UDP-GlcNAc levels in whole worm lysates in the *gfat-1*(*dh783*) R203H mutant points to the predominant role of reduced sensitivity to feedback inhibition as the key gain-of-function mechanism.

R203H GFAT-1 showed reduced sensitivity to UDP-GlcNAc feedback inhibition, indicating a functional role of Arg203 in UDP-GlcNAc inhibition. In wild type GFAT-1, Arg203 is implicated in a salt-bridge network, which might help to position the glutaminase and isomerase domains (Fig. 5f). Previously, we proposed that UDP-GlcNAc promotes a catalytically unfavorable domain orientation, inhibiting GFAT-1^40^. In general, domain movements are common in the family of glutamine amidotransferases and the function of GFAT depends on the relative orientation of the glutaminase and isomerase domains^44^. Mouilleron *et al*. reported a structure of the *E. coli* GFAT active site mutant C1A (PDB 3OOJ) with a drastically changed orientation of the glutaminase domain relative to the isomerase domain, where one monomer adopts an inactive orientation without forming the ammonia channel^45^. This structure underpins the high flexibility of the two domains relative to each other. In wild type *E. coli* GFAT, the glutaminase domain adopts a specific position relative to the isomerase domain after Frc6P binding^16^. The presence of Frc6P activates the glutaminase function of GFAT more than 100-fold in *E. coli* and 70-fold in *C. albicans* and it is very likely that this substrate-induced activation occurs in the human GFAT as well^18,41^. Also, L-Gln binding induces a rotation of the glutaminase domain relative to the isomerase domain in *E. coli* GFAT^19^. These domain movements upon substrate binding are required to enable specific interactions between the isomerase and glutaminase domains that are necessary for catalysis. It is well-established that phosphorylation can introduce conformational changes within loops or domains through altered electrostatic interactions^46^. A well-studied example for a phosphorylation-induced change in domain orientation is rabbit muscle glycogen phosphorylase, whose domains are shifted by approximately 50° relative to each other after phosphorylation^47,48^. The phosphorylation of GFAT-1 at Ser205 would introduce, at physiological pH, a negatively charged phosphate group, potentially affecting ionic interactions in close proximity. Arg202 of the glutaminase domain and Glu425 of the isomerase domain likely form a salt-bridge between the two domains and this interaction might be affected upon Ser205 phosphorylation (Fig. 5f).

Thermofluor assays of wild type and phospho-mimetic GFAT-1 showed a destabilization of the glutaminase domain in the S205D mutant that indicated an altered interdomain interaction. We propose that Ser205 phosphorylation suppresses UDP-GlcNAc inhibition by stabilization of the glutaminase and isomerase domains in a catalytically productive orientation (Fig. 5g). While this conformation does not permit the high catalytic rate of wild type GFAT-1, the fixed conformation of the domains would prevent a UDP-GlcNAc induced domain orientation that might be needed for inhibition.

Our data reveal a UDP-GlcNAc concentration-dependent effect on the activity of GFAT-1 that is phosphorylated at Ser205. This finding explains the previous contradicting reports about the effect of the Ser205 phosphorylation for GFAT-1: Zhou *et al*. observed an activation^32^, while Chang *et al*. reported an inhibition^33^. In both studies, cells were treated with forskolin to activate PKA and the GFAT-1 activity was analyzed in cell lysates. These lysates contained unknown UDP-GlcNAc concentrations that might have affected the results. Moreover, Chang *et al*. confirmed the inhibitory effect *in vitro* with GST-tagged GFAT-1 in the absence of UDP-GlcNAc. However, the data obtained from a GST-tagged GFAT-1 should be taken with caution, because tagging GFAT at the N- or C-terminus interferes with the catalytic reactions^33,49,50^.

Very few enzymes are known in which phosphorylation interferes with feedback inhibition^51^. This uncoupling of the feedback loop by GFAT-1 phosphorylation is an elegant mechanism to rapidly modulate its enzymatic activity after a stress-induced signal. The cAMP-PKA signaling pathway acts downstream of G protein-coupled receptors (GPCR) to mediate signals of neurotransmitters and hormones, such as glucagon or adrenaline^52–55^. The downstream protein targets of the cAMP-PKA signaling pathway regulate glucose homeostasis by inhibition of glycolysis and glycogen synthesis, as well as by triggering glucose release through stimulation of glycogenolysis and gluconeogenesis^55^. Our previous data indicated that GFAT-1 is under a constant UDP-GlcNAc inhibition *in vivo*^40^. Presumably, uncoupling of the feedback inhibition maintains GFAT-1 activity and ensures steady UDP-GlcNAc production when PKA activity is high. Given that UDP-GlcNAc is an essential building block and precursor for all glycosylation reactions in mammals, this mechanism is optimally positioned to ensure a constant UDP-GlcNAc supply. In all, our findings illuminate how the different means of GFAT-1 regulation, kinase signaling, and metabolic feedback, are coordinated at the molecular level to fine tune metabolic flux in the HP.

## Methods

### *C. elegans* strains and culture

All *C. elegans* strains were maintained at 20°C on nematode growth medium (NGM) agar plates seeded with the *E. coli* strain OP50^56^. To provide an isogenic background, the mutant strain was outcrossed against the wild type Bristol N2 strain. The strains used in this study are: N2 (Bristol), *gfat-1*(*dh783*).

### Developmental tunicamycin resistance assay

Gravid adult nematodes were bleached to obtain a synchronized population of eggs, which were transferred to NGM plates containing 10 mg/mL tunicamycin (Sigma-Aldrich) seeded with freeze-killed OP50 *E. coli*. Freeze-killed OP50 bacteria were obtained by three cycles of snap-freezing and thawing of pelleted overnight OP50 bacterial cultures. Worms were kept at 20°C and after four days, healthy day 1 adults were scored, whereas sick larvae were not counted. Throughout the experiment, strain identity was unknown to researchers. Data were assembled upon completion of the experiment.

### Mutant Hawaiian SNP mapping and sequence analysis

Genomic DNA was prepared using the QIAGEN Gentra Puregene Kit according to the manufacturer’s protocol. Whole genome sequencing was conducted on the Illumina HiSeq2000 platform. Paired-end 100 bp reads were used; the average coverage was larger than 16-fold. Sequencing outputs were analyzed using the CloudMap Hawaiian and Variant Discovery Mapping on Hawaiian Mapped Samples (and Variant Calling) Workflow_2-7-2014 pipeline on Galaxy^42,57^. The WS220/ce10 *C. elegans* assembly was used as reference genome.

### Small molecule LC/MS/MS Analysis

UDP-HexNAc concentrations were measured as described previously^39^. In brief, *C. elegans* or HEK293 cells were lysed in water by freeze/thaw cycles, and subjected to chloroform/methanol extraction. Absolute UDP-HexNAc levels were determined using an Acquity UPLC connected to a Xevo TQ Mass Spectrometer (both Waters) and normalized to total protein content.

### Site-directed mutagenesis

A pFL vector for the generation of baculoviruses for the expression of human GFAT-1 isoform 2 (hGFAT-1) with internal His6-tag between Gly299 and Asp300 was cloned previously^40^. The mutations R203H, S205D, and L405R were introduced into pFL-hGFAT1-His299 by site-directed mutagenesis as described previously^58^ (primers: hGFAT1_R203H_for CAAGGCacGGTAGCCCTCTGTTGATTGG, hGFAT1_R203H_rev GAGGGCTACCgtGCCTTGTGCCAACTG, hGFAT1_S205D_for CAAGGCGAGGTgaCCCTCTGTTGATTGG, hGFAT1_S205D_rev GAGGGtcACCTCGCCTTGTGCCAACTG, hGFAT1_L405R_for GTGACTTCCgtGACAGAAACACACCAG, hGFAT1_L405R_rev GTGTTTCTGTCacGGAAGTCACTTGCTAG).

### Baculovirus generation and insect cell expression of full-length GFAT-1

*Sf21* (DSMZ no. ACC 119) suspension cultures were maintained in SFM4Insect™ HyClone™ medium with glutamine (GE Lifesciences) in shaker flasks at 27 °C and 90 rpm in an orbital shaker. GFAT-1 variants were expressed in *Sf21* cells using the MultiBac baculovirus expression system^59^. In brief, hGFAT-1 variants (from the pFL vector) were integrated into the baculovirus genome via Tn7 transposition and maintained as bacterial artificial chromosome in DH10EMBacY *E. coli* cells. Recombinant baculoviruses were generated by transfection of *Sf21* with bacmid DNA. The obtained baculoviruses were used to induce protein expression in *Sf21* cells.

### Bacterial expression of GFAT-1 isomerase domain

The isomerase domain of human GFAT-1 isoform 2 (residues 316-681) was integrated in the plasmid pET28a(+) using NdeI and HindIII restriction sites (primers: hGFAT1-ISO_NdeI_FOR gagCATATGatcatgaagggcaacttcagttcat ttatgc, hGFAT1_HindIII_REV gagAAGCTTtcactctacagtcacagatttggca agattc). This vector was used to recombinantly express the isomerase domain with N- terminal His6tag and a thrombin cleavage site under the control of the T7 promoter in Rosetta (DE3) *E. coli*. LB cultures were incubated at 37°C and 180 rpm until an OD_600_ of 0.4-0.6 was reached. Then, protein expression was induced by addition of 0.5 mM isopropyl-β-D-1-thiogalactopyranosid and incubated for 3 h at 37°C and 180 rpm. Cultures were harvested and pellets stored at −80°C.

### GFAT-1 purification

*Sf21* cells (full-length GFAT-1) or *E. coli* (isomerase domain) were lysed by sonication in lysis buffer (50 mM Tris/HCl pH 7.5, 200 mM NaCl, 10 mM Imidazole, 2 mM Tris(2-carboxyethyl)phosphin (TCEP), 0.5 mM Na2Frc6P, 10% (v/v) glycerol, supplemented with complete EDTA-free protease inhibitor cocktail (Roche) and 10 μg/ml DNAseI (Sigma)). Cell debris and protein aggregates were removed by centrifugation and the supernatant was loaded on a Ni-NTA Superflow affinity resin (Qiagen). The resin was washed with lysis buffer and the protein eluted with lysis buffer containing 200 mM imidazole. For the isomerase domain, the His6-tag was proteolytically removed using 5 U of thrombin (Sigma-Aldrich) per mg protein overnight at 4°C. The isomerase domain was again purified by IMAC in order to remove undigested His6-tagged protein. Full-length GFAT-1 and the isomerase domain were further purified according to their size on a HiLoad^™^ 16/60 Superdex™ 200 prep grade prepacked column (GE Healthcare) using an ÄKTAprime chromatography system at 4°C with a SEC buffer containing 50 mM Tris/HCl, pH 7.5, 2 mM TCEP, 0.5 mM Na2Frc6P, and 10% (v/v) glycerol.

### Crystallization

For crystallization experiments, the SEC buffer was supplemented with 50 mM L-Arg and 50 mM L-Glu to improve protein solubility^60^. GFAT-1 was crystallized at a concentration of 8 mg/ml in sitting-drops by vapor diffusion at 20°C. Crystals grew in the PACT *premier™* HT-96 (Molecular Dimensions) screen with a reservoir solution containing 0.1 M Bis tris propane pH 8.5, 0.2 M Potassium sodium tartrate and 20% (w/v) PEG3350 and were further optimized. The optimization screen was set up with drop ratios of 1.5 μl protein solution to 1.5 μl precipitant solution and 2 μl protein solution to 1 μl precipitant solution. Best crystals grew in a broad range of 0.1 M Bis tris propane pH 8.5 to 9.0, 0.2 to 0.4 M potassium sodium tartrate, and 20 % (w/v) PEG3350. Data were collected from crystals cryoprotected with reservoir solution supplemented with 15% (v/v) glycerol.

### Data collection and refinement

X-ray diffraction measurements were performed at beamline X06DA at the Swiss Light Source, Paul Scherrer Institute, Villigen (Switzerland). The mutant GFAT-1 structures were determined by molecular replacement with phenix.phaser^61,62^ using the model of the wild type human GFAT-1 (PDB 6R4E) as search model. GFAT-1 was further manually built using COOT^63^ and iterative refinement rounds were performed using phenix.refine^62^. Geometry restraints for ligands were generated with phenix.elbow software^62^. Structures were visualized using PyMOL (Schrödinger).

### GDH-coupled activity assay and UDP-GlcNAc inhibition

GFAT’s amidohydrolysis activity was measured with a coupled enzymatic assay using bovine glutamate dehydrogenase (GDH, Sigma Aldrich G2626) in 96 well standard microplates (F-bottom, BRAND #781602) as previously described^50^ with small modifications. In brief, the reaction mixtures contained 6 mM Frc6P, 1 mM APAD, 1 mM EDTA, 50 mM KCl, 100 mM potassium-phosphate buffer pH 7.5, 6.5 U GDH per 96 well and for L-Gln kinetics varying concentrations of L-Gln. For UDP-GlcNAc inhibition assays the L-Gln concentration was kept at 10 mM. The plate was pre-warmed at 37°C for 10 min and the activity after enzyme addition was monitored continuously at 363 nm in a microplate reader. The amount of formed APADH was calculated with ε_(363 nm, APADH)_ = 9100 l*mol^-1^*cm^-1^. Reaction rates were determined by Excel (Microsoft) and Km, vmax, and IC_50_ were obtained from Michaelis Menten or dose response curves, which were fitted by Prism 7 or 8 software (Graphpad).

### GNA-1 expression and purification

The expression plasmid for human GNA-1 with N-terminal His6-tag was cloned previously^40^. Human GNA-1 with N-terminal His6-tag was expressed in Rosetta (DE3) *E. coli* cells. LB cultures were incubated at 37°C and 180 rpm until an OD600 of 0.4-0.6 was reached. Then, protein expression was induced by addition of 0.5 mM isopropyl-β-D-1-thiogalactopyranosid and incubated for 3 h at 37°C and 180 rpm. Cultures were harvested and pellets stored at −80°C. Human GNA-1 purification protocol was adopted from Hurtado-Guerrero et al.^64^ with small modifications. *E. coli* were lysed in 50 mM HEPES/NaOH pH 7.2, 500 mM NaCl, 10 mM imidazole, 2 mM 2-mercaptoethanol, 5% (v/v) glycerol with complete EDTA-free protease inhibitor cocktail (Roche) and 10 μg/ml DNAseI (Sigma) by sonication. The lysate was clarified by centrifugation and the supernatant loaded on Ni-NTA Superflow affinity resin (Qiagen). The resin was washed with wash buffer (50 mM HEPES/NaOH pH 7.2, 500 mM NaCl, 50 mM imidazole, 5% (v/v) glycerol) and the protein was eluted with wash buffer containing 250 mM imidazole. Eluted protein was then dialyzed against storage buffer (20 mM HEPES/NaOH pH 7.2, 500 mM NaCl, 5% (v/v) glycerol).

### GNA-1 and GNA-1-coupled activity assays

The activity of human GNA-1 was measured in 96 well standard microplates (F-bottom, BRAND #781602) as described previously^65^. For kinetic measurements, the assay mixture contained 0.5 mM Ac-CoA, 0.5 mM DTNB, 1 mM EDTA, 50 mM Tris/HCl pH 7.5 and varying concentrations of D-GlcN6P. The plates were pre-warmed at 37°C and reactions were initiated by addition of GNA-1. The absorbance at 412 nm was followed continuously at 37°C in a microplate reader. The amount of produced TNB, which matches CoA production, was calculated with ε_(412 nm, TNB)_ = 13800 l*mol^-1^*cm^-1^. Typically, GNA-1 preparations showed a K_m_ of 0.2 ± 0.1 mM and a kcat of 41 ± 8 sec^-1^.

GFAT’s D-GlcN6P production was measured in a GNA-1-coupled activity assay following the consumption of AcCoA at 230 nm in UV transparent 96 well microplates (F-bottom, Brand #781614) as described by Li et al.^65^. In brief, the assay mixture contained 10 mM L-Gln, 0.1 mM AcCoA, 50 mM Tris/HCl pH 7.5, 2 μg hGNA-1 and varying concentrations of Frc6P. The plates were incubated at 37°C for 4 min and reactions started by adding L-Gln. Activity was monitored continuously at 230 nm and 37°C in a microplate reader. The amount of consumed AcCoA was calculated with ε_(230 nm, AcCoA)_ = 6436 l*mol^-1^*cm^-1^. As UDP-GlcNAc absorbs light at 230 nm, the GNA-1-coupled assay cannot be used to analyze UDP-GlcNAc effects on activity.

### Alignments

The protein sequence alignments were created with following UnitProt IDs: *Caenorhabditis elegans:* Q95QM8, *Homo sapiens* isoform 2: Q06210-2, *Mus musculus* isoform 2: P47856-2, *Candida albicans:* P53704, *Escherichia coli:* P17169. The alignment of *H. sapiens* and *C. elegans* GFAT was formatted with the ESPript3 server (*espript.ibcp.fr/*)^66^.

For the structural superposition, the wild type human GFAT-1 structure in complex with L-Glu and Glc6P (PDB ID 6R4E) was aligned with the structures of mutant GFAT-1 R203H and L405R. UDP-GlcNAc was displayed in the structures after superposition with the structure of wild type human GFAT-1 in complex with L-Glu, Glc6P, and UDP-GlcNAc (PDB ID 6SVP).

### *In vitro* PKA phosphorylation for Protein Mass Spectrometry

Protein Kinase A Catalytic Subunit from bovine heart (PKA, EC 2.7.11.11, Sigma) was reconstituted in bi-distilled water containing 6 mg/ml DTT at a concentration of 50 μg/ml. Purified GFAT-1 variants were phosphorylated in an assay mixture containing 10 mM MgCl2, 2 mM Na-ATP, and 20 U PKA in 100 μl GFAT-1 SEC buffer (50 mM Tris/HCl, pH 7.5, 2 mM TCEP, 0.5 mM Na2Frc6P, and 10% (v/v) glycerol) for 30 min at 30°C. The samples were stored frozen. For proteomic analysis, 5 μg GFAT-1 was alkylated by 5 mM chloroacetamide, reduced with 1 mM TCEP, and digested by 0.1 μg Lys-C Endoproteinase (MS Grade, ThermoScientific) in 50 mM Tris/HCl, pH 8.3 overnight in a waterbath at 37°C. The digest was acidified by addition of formic acid (end concentration 0.1%) and the resulting peptides were purified using C-18 STAGE (STop And Go Extraction) tips^67^.

### Untargeted protein Mass Spectrometry

One fifth of the desalted STAGE tip purified peptides were separated on a 25 cm, 75 μm internal diameter PicoFrit analytical column (New Objective) packed with 1.9 μm ReproSil-Pur 120 C18-AQ media (Dr. Maisch HPLC GmbH) using an EASY-nLC 1200 (Thermo Fisher Scientific). The column was maintained at 50°C. Buffer A and B were 0.1% formic acid in water and 0.1% formic acid in 80% acetonitrile. Peptides were separated on a segmented gradient from 6% to 31% buffer B for 57 min and from 31% to 44% buffer B for 5 min at 200 nl/min. Eluting peptides were analyzed on an Orbitrap Fusion Tribrid mass spectrometer (Thermo Fisher Scientific). Peptide precursor m/z was measured at 60000 resolution in the 350 to 1500 m/z range. Precursors with charge state from 2 to 7 only were selected for HCD fragmentation using 27% normalized collision energy. The m/z values of the peptide fragments were measured at a resolution of 30000 using an AGC target of 2e5 and 80 ms maximum injection time. Upon fragmentation, precursors were put on a dynamic exclusion list for 45 sec.

Heavy synthetic peptides, corresponding to phosphorylated Ser205, read below, were separated on a segmented gradient from 6% to 60% buffer B for 62 min at 200 nl/min. The mass spectrometric analysis was carried out as described above.

Raw data were analyzed with MaxQuant version 1.6.1.0 using the integrated Andromeda search engine^68,69^. Peptide fragmentation spectra were searched against manually created GFAT-1 fasta file. The database was automatically complemented with sequences of contaminating proteins by MaxQuant. Methionine oxidation, protein N-terminal acetylation and Phospho (STY) were set as variable modifications; cysteine carbamidomethylation was set as fixed modification. The digestion parameters were set to “specific” and “LysC/P,” The minimum number of peptides and razor peptides for protein identification was 1; the minimum number of unique peptides was 0. Protein identification was performed at a peptide spectrum matches and protein false discovery rate of 0.01. For the analysis of the raw data from the synthetic heave peptides, Lys8 was set as “Standard” and used as a variable modification.

### Targeted protein Mass Spectrometry

The phosphorylated (Ser205, pS), heavily labeled (K*) reference peptides Maxi SpikeTides L (wild type: SVHFPGQAVGTRRG-pS-PLLIGVRSEH-K* and mutant R203H: SVHFPGQAVGTRHG-pS-PLLIGVRSEH-K*) were purchased from JPT Peptide Technologies (Berlin, Germany). Heavy peptides were dissolved in 50% acetonitrile, 0.05% formic acid in water and sonicated in a water bath for one minute. The solution was further diluted using 0.1% formic acid in water for a final peptide concentration of 7 nM. The solution was aliquoted and stored at −20°C. For analysis, desalted GFAT-1 peptides were dissolved in 10 μl 0.1% formic acid in water and 2 μl of the peptide solution were combined with 2 μl of the tenfold diluted heavy peptide solution; 2 μl were analyzed by targeted mass spectrometry.

For targeted analysis, peptides were separated as described above. Targeted analysis was performed on an Orbitrap Fusion Tribrid mass spectrometer (Thermo Fisher Scientific). The m/z values for charge states 4 and 5 from the heavy and light peptide were chosen for targeted fragmentation across the entire LC run. The isolation width was 1.6 and collision energy was set to 27%. Fragment m/z values were measured in the Orbitrap in profile mode, at a resolution of 60K, an AGC target of 5e4, and a maximum injection time of 118 ms.

Raw data was analyzed using Skyline^70^ version 19.1.0.193. A library was built from the untargeted analysis of the heavy peptides. The library ion match tolerance was set to 0.05 m/z, the 10 most intense product ions were picked. Ions with charges 1 or 2 and type y an b were used for quantification. An isotope modification of type “Label” was created using the following modification: 13C(6)25N(2) (K). Isotope label type and internal standard type were set to “heavy”. For quantification, MS level was set to 2. Results at the peptide and the precursor level were exported and used for downstream data analysis and visualization.

### Thermofluor assay

The thermal stability of proteins was analyzed by thermofluor assays. For that purpose, the proteins were incubated with SYPRO orange dye (Sigma-Aldrich), which binds specifically to hydrophobic amino acids leading to an increased fluorescence at 610 nm when excited with a wavelength of 490 nm. The melting temperature is defined as the midpoint of temperature of the protein-unfolding transition^71^. This turning point of the melting curve was extracted from the derivative values of the RFU curve, where a turning point to the right is a minimum. The thermal stability of the isomerase domain of hGFAT-1 and hGFAT-1 variants, were determined in SEC buffer. The influence of increasing salt concentrations (0 to 1 M) in the SEC buffer was assessed for the isomerase domain of hGFAT-1 and full-length hGFAT-1. The reaction mixtures were pipetted in white RT-PCR plates and contained 5 μl SYPRO orange dye (1:500 dilution in ddH2O) and 5-10 μg protein in a total volume of 50 μl. The plates were closed with optically clear tape and placed in a BioRad CFX-96 Real-Time PCR machine. The melting curves were measured at 1°C/min at the FRET channel and the data analyzed with CFX Manager™ (BioRad).

### Circular dichroism spectroscopy

For CD measurements, GFAT-1 was dialyzed in 10 mM potassium phosphate buffer pH 7.5, 0.5 mM TCEP, 0.5 mM Na2Frc6P, and 10% (v/v) glycerol, and the protein concentration was adjusted to 0.2 mg/ml. The UV spectra in the range of 195–260 nm were recorded with a J-715 CD spectropolarimeter (Jasco, Gross-Umstadt, Germany) at 20°C using a quartz cuvette with 0.1 cm path length. The buffer baseline was recorded separately and subtracted from each sample spectrum. The obtained ellipticity (θ, deg) was converted to mean residue ellipticity [θ] using: [θ] = θ/(10*n*c*l) in deg*cm^2^*dmol^-1^ (n is the number of amino acids, c the protein concentration in mol/l, and l the path length of the cuvette).

### Mammalian cell culture and stable cell line generation

HEK293 cells (ATCC) were cultured at 37°C, 5% CO_2_ on treated polystyrene culture dishes (Corning) in DMEM media with high glucose (4.5 g/l; 25 mM) with pyruvate (Gibco, 11995-065) supplemented with 10% fetal bovine serum (Gibco), 100 U/ml penicillin and 100 μg/ml streptomycin (Gibco).

Cell lines stably overexpressing GFAT-1 variants were generated by transfection of HEK293 cells with pcDNA3.1 plasmids with human GFAT-1 containing an internal His6-tag between Gly299 and Asp300 (pcDNA3.1-hGFAT-1-His299). Internally tagged GFAT-1 was subcloned from pFL-hGFAT1-His299 using NheI and HindIII restriction sites. For each variant, one well of a 6 well plate was transfected with 2 μg of plasmid DNA with Lipofectamine^®^ 2000 (Life Technologies™) according to the manufacturer’s protocol. The selection was performed with 500 μg/ml G418 for several weeks. For Western blot analysis, cells were washed, collected and the proteins from the pellets were extracted by lysis in RIPA-buffer (120 mM NaCl, 50 mM Tris/HCl, 1% (v/v) NP40, 0.5% (w/v) deoxycholate, 0.1% (w/v) SDS; pH 7.5). The cell debris was removed by centrifugation and the protein concentration of the supernatants were determined using Pierce™ bicinchoninic acid (BCA) protein assay kit (Thermo Scientific) according to manufacturer’s protocol. 2 μg cell lysate was separated by reducing SDS-PAGE and transferred to PVDF membranes. Primary antibodies against human GFAT1 and a-TUBULIN were used. Chemiluminescence of the appropriate secondary rabbit or mouse HRP-conjugated antibodies after incubation with ECL HRP substrate (Immobilon^™^ Western HRP Substrate, Millipore) was detected using a Chemi Doc^™^ Quantity One^®^ system (Bio-Rad).

### Antibodies

The following antibodies were used in this study: GFAT1 (rb, EPR4854, Abcam ab125069, 1:1000), a-TUBULIN (ms, DM1A, Sigma T6199, 1:50000, rabbit IgG (gt, LifeTechnologies G21234, 1:5000), and mouse IgG (gt, LifeTechnologies G21040, 1:5000).

## Data availability

Structural data reported in this study have been deposited in the Protein Data Bank with the accession codes 6ZMJ and 6ZMK. The mass spectrometry proteomics data have been deposited to the ProteomeXchange Consortium via the PRIDE^72^ partner repository with the dataset identifier PXD020451. All other data supporting the presented findings are available from the corresponding authors upon request.

## Acknowledgments

We thank all M.S.D. and U.B. laboratory members for helpful discussions. The *gfat-1(dh783) C. elegans* strain was kindly provided by Adam Antebi (Max Planck Institute for Biology of Ageing). Whole genome sequencing was done at the Cologne Center for Genomics. We thank Yvonne Hinze and Patrick Giavalisco from the metabolomics core facility of the Max Planck Institute for Biology of Ageing. We are grateful to Schirin Birkmann for support in the insect cell maintenance. Crystals were grown in the Cologne Crystallization facility (C2f). We thank the staff of beamline X06DA at the Swiss Light Source, Paul Scherrer Institute, Villigen (Switzerland) for their support during data collection. This work was supported by the German Federal Ministry of Education and Research (BMBF, grant 01GQ1423A EndoProtect), by the German Research Foundation (DFG, Projektnummer 73111208-SFB 829, B11 and B14), by the European Commission (ERC-2014-StG-640254-MetAGEn), and by the Max Planck Society. The Cologne Crystallization Facility C2f was supported by DFG grant INST 216/949-1 FUGG.

## Author contributions

S.R. and M.S.D. designed the project. S.R. performed the biochemical and crystallization experiments, as well as the cell culture experiments. F.M. performed the experiments related to *C. elegans*. S.M. helped with the mammalian cell culture experiments. I.A. did the protein mass spectrometry measurements and analysis. S.R., M.S.D., and U.B. wrote the manuscript. S.R. prepared the figures.

## Competing interests

The authors declare no competing interest.

## Supplementary figures

**Fig. 1 Supp.:**
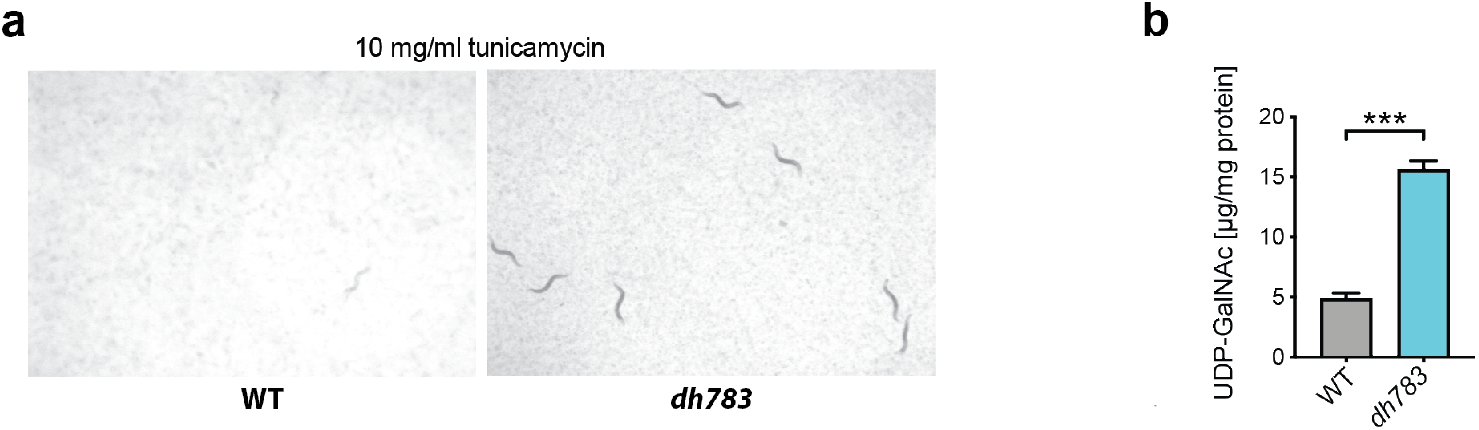
Characterization of *gfat-1(dh783) C. elegans* mutants. **a,** Representative images of wild type (N2) and *gfat-1*(*dh783*) animals 4 days post-hatch on NGM plates containing 10 mg/ml tunicamycin. **b,** UDP-GalNAc levels in N2 wild type and *gfat-1*(*dh783*) animals (mean +SD, n=5, *** p<0.001, unpaired t-test).

**Fig. 2 Supp:**
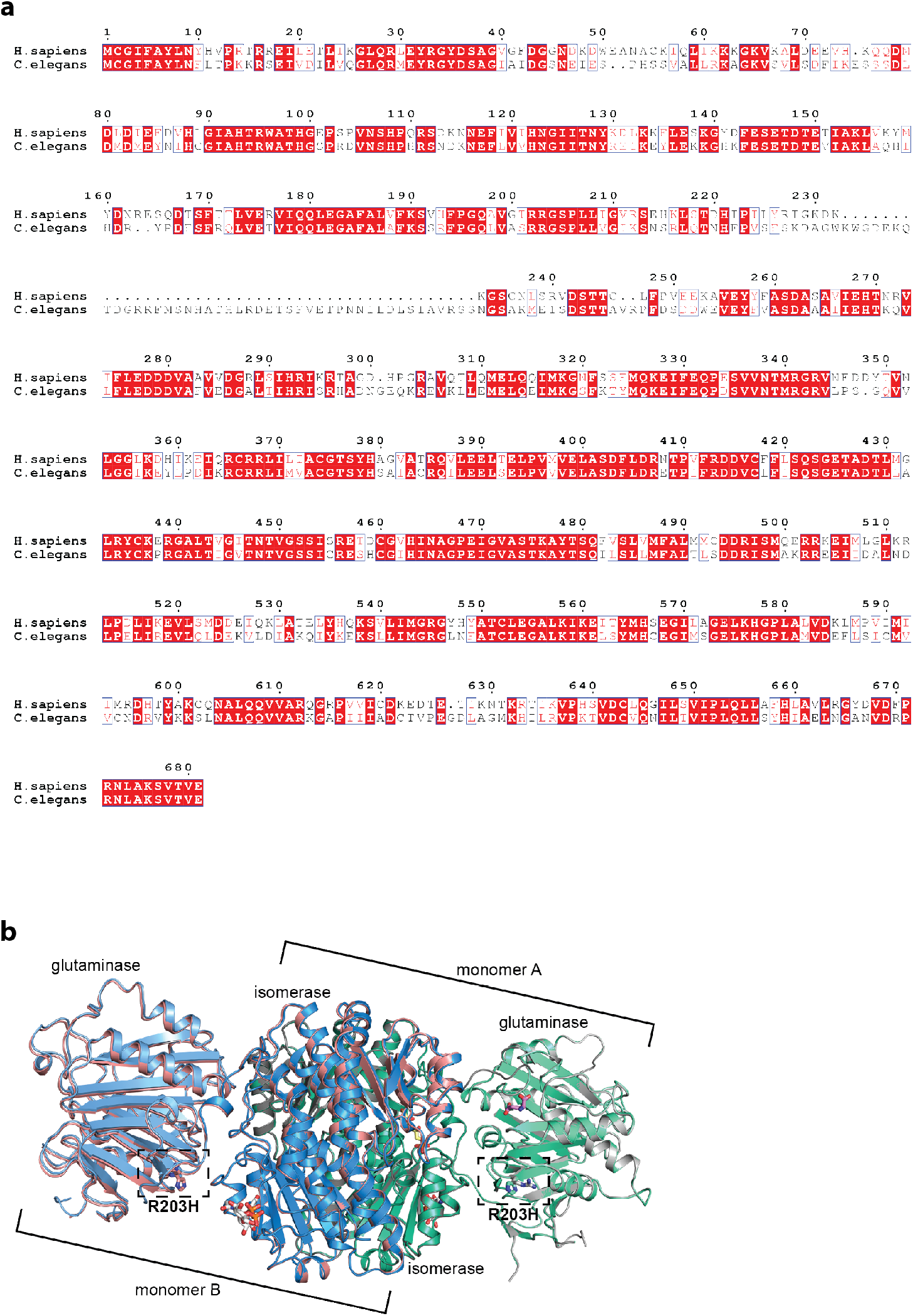
The GFAT-1 R203H gain-of-function substitution perturbs UDP-GlcNAc feedback inhibition. **a**, Protein sequence alignment of GFAT-1. Red boxes indicate identical residues, red letters indicate similar residues. **b,** Position of the R203H mutation in the dimeric structure of GFAT-1. Proteins are presented as cartoons. Superposition of wild type GFAT-1 (light gray/dark gray) and R203H GFAT-1 (green-cyan, teal). Glc6P (yellow sticks), L-Glu (violet sticks), and UDP-GlcNAc (white sticks) are highlighted, as well as the position of R203H (black box).

**Fig. 3 Supp.:**
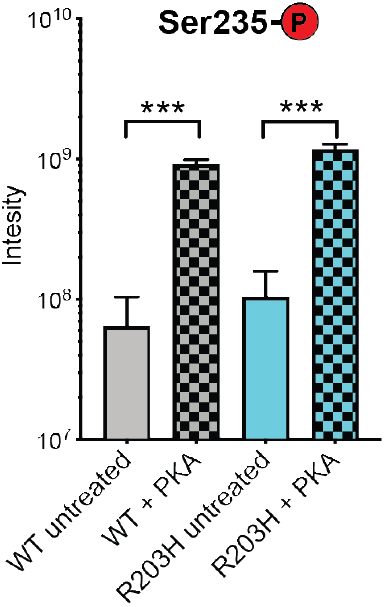
*In vitro* PKA treatment of GFAT-1 results in phosphorylation at Ser235. Intensity of phosphorylated peptides at Ser235 of wild type GFAT-1 (gray) and R203H (cyan) before and after treatment with PKA normalized to protein abundance (mean +SEM, n=4, *** p<0.001, one-way ANOVA).

**Fig. 4 Supp:**
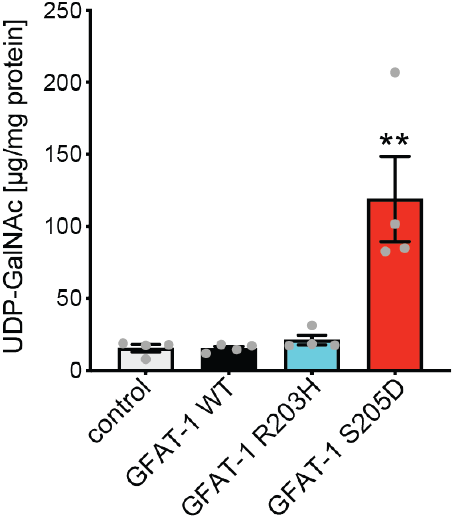
PKA phosphorylation at Ser205 modulates UDP-GlcNAc inhibition of GFAT-1. LC/MS measurement of UDP-GalNAc normalized to protein content presented as means +SEM with n=4, ** p<0.01 versus WT, one-way ANOVA.

**Fig. 5 Supp:**
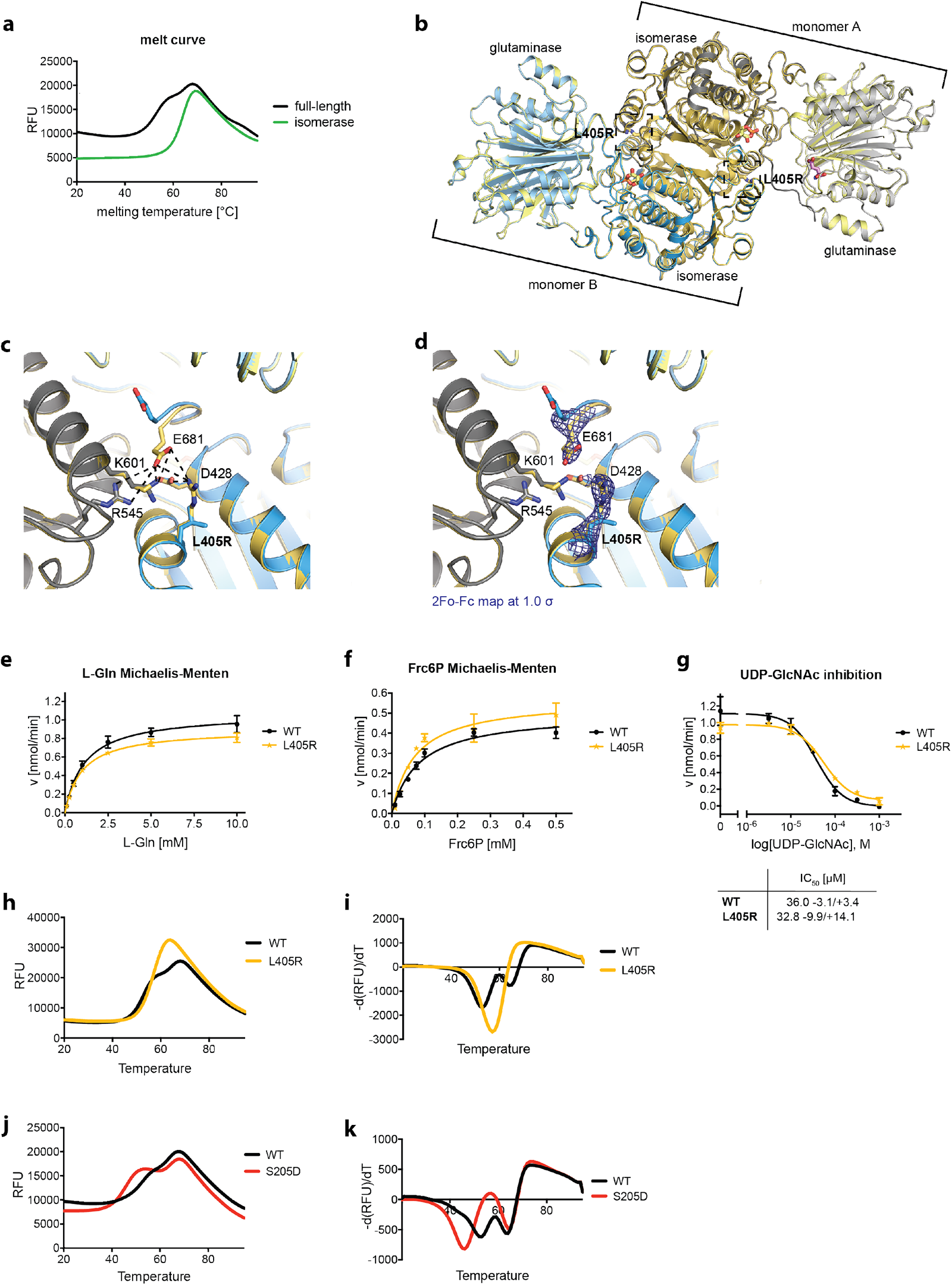
Altered domain stability of GFAT-1 after PKA phosphorylation at Ser205. **a**, Representative melting curves of full-length (black) and isolated isomerase domain (green) in standard SEC buffer without NaCl. **b-d**, Position of the L405R mutation in the structure of GFAT-1. Superposition of the structures of wild type (monomer A gray, monomer B blue) and L405R (yellow) GFAT-1 in cartoon representation. **b**, Overview of the dimers. Glc6P (yellow sticks) and L-Glu (violet sticks) are highlighted, as well as the mutation L405R (black boxes). **c**, Close-up of the interactions of L405R. **d**, Close-up of the electron density of L405R and Glu681. The 2Fo-Fc map of Arg405 and Glu681 are colored dark blue and their contour levels are at 1.5 RMSD. **e**, L-Gln kinetics of wild type (black circles) and L405R (yellow stars) GFAT-1 (mean ±SEM, WT n=5, L40R n=3). **f**, Frc6P kinetics of wild type (black circles) and L405R (yellow stars) GFAT-1 (mean ±SEM, WT n=5, L405R n=3). **g**, Representative UDP-GlcNAc dose response assay of wild type (black circles) and L405R (yellow stars) GFAT-1 (mean ±SD, n=3). Table: IC_50_ UDP-GlcNAc values (mean ±SEM, n=3). **h**, Representative melting curves of wild type (black) and L405R (yellow). **i**, Representative derivative melting curves of wild type (black) and L405R (yellow). **j**, Representative melting curves of wild type (black) and S205D (red). **k**, Representative derivative melting curves of wild type (black) and S205D (red).

## Notes

https://www.ebi.ac.uk/pride/archive/projects/PXD020451

### Competing Interest Statement

The authors have declared no competing interest.

